# The Retrieval-related Anterior shift is Moderated by Age and Correlates with Memory Performance

**DOI:** 10.1101/2021.08.30.457871

**Authors:** Sabina Srokova, Paul F. Hill, Michael D. Rugg

**Affiliations:** Center for Vital Longevity, University of Texas at Dallas, Dallas, TX 75235; School of Behavioral and Brain Sciences, University of Texas at Dallas, Richardson, TX 75080; Department of Psychology, University of Arizona, Tucson, AZ 85721; School of Psychology, University of East Anglia, Norwich NR4 7TJ, United Kingdom

## Abstract

Recent research suggests that episodic memory is associated with systematic differences in the localization of neural activity observed during memory encoding and retrieval. The retrieval-related anterior shift is a phenomenon whereby the retrieval of a stimulus event (e.g., a scene image) is associated with a peak neural response which is localized more anteriorly than the response elicited when the stimulus is experienced directly. Here, we examine whether the magnitude of the anterior shift, i.e., the distance between encoding- and retrieval-related response peaks, is moderated by age, and also whether the shift is associated with memory performance. Younger and older human subjects of both sexes underwent fMRI as they completed encoding and retrieval tasks on word-face and word-scene pairs. We localized peak scene- and face-selectivity for each individual participant within the face-selective precuneus (PCU) and in three scene-selective (parahippocampal place area [PPA], medial place area [MPA], occipital place area [OPA]) regions of interest (ROIs). In line with recent findings, we identified an anterior shift in PPA and OPA in both age groups and, in older adults only, in MPA and PCU also. Of importance, the magnitude of the anterior shift was larger in older than in younger adults. The shift within the PPA exhibited an age-invariant across-participant negative correlation with source memory performance, such that a smaller displacement between encoding- and retrieval-related neural activity was associated with better performance. These findings provide novel insights into the functional significance of the anterior shift, especially in relation to memory decline in older age.

**Significance Statement:** Cognitive aging is associated with reduced ability to retrieve precise details of previously experienced events. The retrieval-related anterior shift is a phenomenon in which category-selective cortical activity at retrieval is localized anterior to the peak activity at encoding. The shift is thought to reflect a bias at retrieval in favor of semantic and abstract information at the expense of low-level perceptual detail. Here, we report that the anterior shift is exaggerated in older relative to younger adults, and we demonstrate that a large shift in the parahippocampal place area is associated with poorer memory performance. These findings suggest that the shift is sensitive to increasing age and that it is moderated by the quality and content of the retrieved episode.

## 1. Introduction

Cognitive aging is associated with a disproportionate decline in memory for contextual details of previously experienced episodes. Relative to younger adults, older adults tend to retrieve memories with less specificity and fewer details (Levine et al., 2002; Addis et al., 2008), while memory for semantic information and general knowledge remains relatively preserved (Nilsson, 2003; Nyberg et al., 2012). Recent findings suggest that even when a memory of an event is successfully retrieved, the precision and specificity of the retrieved content is reduced with increasing age (Nilakantan et al., 2018; Korkki et al., 2020). These findings are consistent with the notion that older adults rely on relatively abstract or ‘gist-like’ memories and experience a reduction in episodic detail (Koustaal & Schacter, 1997; Dennis et al., 2007; 2008, Gallo et al., 2019).

Episodic memory retrieval is associated with the ‘reactivation’ of patterns of cortical activity elicited when the episode was experienced, a phenomenon termed cortical reinstatement (for reviews see Danker and Anderson, 2010; Rissman and Wagner, 2012; Rugg et al., 2015; Xue, 2018). The strength and specificity of cortical reinstatement have been reported to be reduced in older age (Bowman et al., 2019; St-Laurent & Buchsbaum, 2019; Folville et al., 2020; Hill et al., 2021, but see Wang et al., 2016; Thakral et al., 2017). The strength of cortical reinstatement has also been reported to predict the likelihood of successful retrieval, leading to the proposal that cortical reinstatement indexes the amount of retrieved episodic content (e.g., Johnson et al., 2009; Trelle et al., 2020; Hill et al., 2021). Thus, age-related reductions in the strength of cortical reinstatement may reflect older adults’ tendency to retrieve less detailed episodic information than their younger counterparts.

Whereas cortical reinstatement is a well-established phenomenon, recent research demonstrates that there are systematic differences in the localization of content-selective cortical activity observed at encoding and retrieval, thus challenging the notion that the neural populations active at encoding are merely reactivated at retrieval. Mental imagery and retrieval of perceptual stimuli (e.g., scene images) have been reported to be associated with neural activation that peaks slightly anterior to the regions maximally recruited during direct perception of the stimuli (for review, see Favilla et al., 2020). This retrieval-related bias towards more anterior neural recruitment has been termed the ‘anterior shift’ (e.g., Rugg & Thompson-Schill, 2013; Bainbridge et al., 2021). The functional significance of this shift is largely unknown, although it has been suggested that it reflects a ‘transformation’ of a mnemonic representation such that different attributes of an event (such as perceptual details) are differentially emphasized at encoding and retrieval. (Favilla et al., 2020). Given that the posterior-anterior axis of occipito-temporal cortex has been held to be hierarchically organized, forming a gradient of increasing abstraction, the anterior shift may reflect a shift towards abstracted representations that emphasize conceptual attributes of a stimulus event at the expense of ‘lower-level’ perceptual and sensory features (e.g., Peelen & Caramazza, 2012; Martin et al., 2018).

Here, younger and older adults underwent fMRI as they viewed concrete words paired with images of faces and scenes. Participants remained in the scanner to complete a retrieval task during which they were presented with old or novel words under the requirement to retrieve the image associated with each word judged to be old. We addressed two key questions. First, we examined whether the anterior shift is moderated by age. In light of evidence suggesting that older adults tend to retrieve more gist-like (abstracted) memories than younger individuals, the aforementioned ‘abstraction’ account of the anterior shift leads to the prediction that it will be exaggerated in older relative to young adults. Second, we asked whether the anterior shift is a moderator of individual differences in memory performance. According to the abstraction account, to the extent that a memory test depends on the retrieval of detailed perceptual information, a negative relationship between the magnitude of the shift and memory performance is predicted.

## 2. Materials and Methods

Outcomes of analyses of data from the present experiment have been described in two prior reports (Srokova et al., 2020; Hill et al., 2021). Descriptions of the experimental design, procedure, and the outcomes of the behavioral analyses were reported previously and are summarized here for the convenience of the reader. The analyses of the retrieval-related ‘anterior shift’ described below have not been reported previously.

All experimental procedures were approved by the Institutional Review Boards of The University of Texas at Dallas and The University of Texas Southwestern Medical Center. Each participant gave informed consent prior to their participation in the study.

### 2.1. Participants

Twenty-seven younger and 33 older adult participants were recruited from the University of Texas at Dallas and surrounding metropolitan Dallas communities. All participants were compensated $30/hour and were reimbursed up to $20 for travel. Participants were right-handed, had normal or corrected-to-normal vision, and were fluent English speakers before the age of five. None of the participants had a history of neurological or cardiovascular disease, diabetes, substance abuse, or current or recent use of prescription medication affecting the central nervous system. Potential MRI-eligible participants were excluded if they did not meet pre-determined performance criteria on our neuropsychological test battery (see below).

Three younger and nine older adults were excluded from subsequent analyses. Two participants voluntarily withdrew from the study and one participant was excluded due to technical difficulties during MRI scanning. Additionally, the behavioral performance of two participants resulted in critical memory bins having too few trials, and six participants were excluded due to at- or near-chance source memory performance (probability of source recollection, pSR, < 0.1). Lastly, one participant was excluded due to an incidental MRI finding. The final sample consisted of 24 younger adults (15 females; age range = 18-28 years, M (SD) = 22.4 (3.2) years) and 24 older adults (14 females; age range = 65-75 years, M (SD) = 70.1 (3.4) years).

### 2.2. Neuropsychological Testing

All participants completed a neuropsychological test battery which was administered on a separate day prior to participation in the fMRI session. The battery consisted of the following tests: Mini-Mental State Examination (MMSE), the California Verbal Learning Test II (CVLT: Delis et al., 2000), Wechsler Logical Memory Tests 1 and 2 (Wechsler, 2009), the Symbol Digit Modalities test (SDMT; Smith, 1982), the Trail Making Tests A and B (Reitan & Wolfson, 1985), the F-A-S subtest of the Neurosensory Center Comprehensive Evaluation for Aphasia (Spreen & Benton, 1977), the Forward and Backward digit span subtests of the revised Wechsler Adult Intelligence Scale (Wechsler, 1981), Category fluency test (Benton, 1968). Raven’s Progressive Matrices List I (Raven et al., 2000), and the Wechsler Test of Adult Reading (WTAR; Wechsler, 2001). Potential participants were not accepted into the study for any of the following reasons: if their MMSE score was below 27, if they scored more than 1.5 standard deviations below age- and education-adjusted norms on one or more long-term memory test or on at least two non-memory tests, or if their estimated full-scale IQ was less than 100. These criteria were employed to minimize the likelihood of including older participants with mild cognitive impairment or early dementia.

### 2.3. Experimental Materials

Experimental stimuli were presented using Cogent 2000 (www.vislab.ucl.ac.uk/cogent_2000.php) implemented in Matlab (www.mathworks.com). The study and test phases were completed inside the scanner, and stimuli were projected onto a translucent screen placed at the rear end of the scanner bore and viewed through a mirror fixed onto the head coil. The critical experimental stimuli comprised 288 concrete nouns, 96 colored images of faces (48 male, 48 female) and 96 colored images of scenes (48 urban, 48 rural). An additional 68 words and 40 images were used as practice stimuli or as filler trials during the experiment proper. The critical stimuli were used to create 24 stimulus lists which were assigned to yoked pairs of younger and older participants. Each study list consisted of 192 randomly selected word-image pairs interspersed with 96 null trials (white fixation cross) and divided into 4 study blocks. Consequently, a single study block comprised 48 critical word-image trials (divided equally between male and female faces, and urban and rural scenes) and 24 null trials. The test list comprised 192 old (studied) trials, 96 new trials, and 96 null trials, evenly distributed into 4 test blocks. The orderings of the items in the study and test lists were pseudorandomized while ensuring that participants experienced no more than three consecutive critical trials of the same image category, no more than three new trials, and no more than two null trials.

### 2.4. Experimental Procedure

Participants received instructions and completed practice study and test tasks prior to entering the scanner. Participants then underwent fMRI as they completed two study-test cycles. Each cycle consisted of two study runs (approx. 8 minutes each) followed by two test runs (approx. 10 minutes each). A schematic of the study and test tasks is illustrated in Figure 1. Each study trial began with a red fixation cross presented for 500 ms. The fixation cross was followed by the presentation of the word-image pair, which remained on the screen for 2000ms. When presented with an image of a face, participants were to imagine the person in the image interacting with the object. During scene trials, participants imagined the object interacting or moving around within the scene. Participants rated the vividness of the imagined scenario on a 3-point scale (“not vivid”, “somewhat vivid”, “very vivid”) using a scanner-compatible button box and the index, middle, and ring fingers of the right hand, respectively. The presentation of the word-image pair was followed by a white fixation cross that lasted for an additional 2000 ms. Participants were allowed to make their vividness response from the onset of the word-image pair until the termination of the white fixation cross, thus providing a 4000 ms response window.

**Figure 1:**
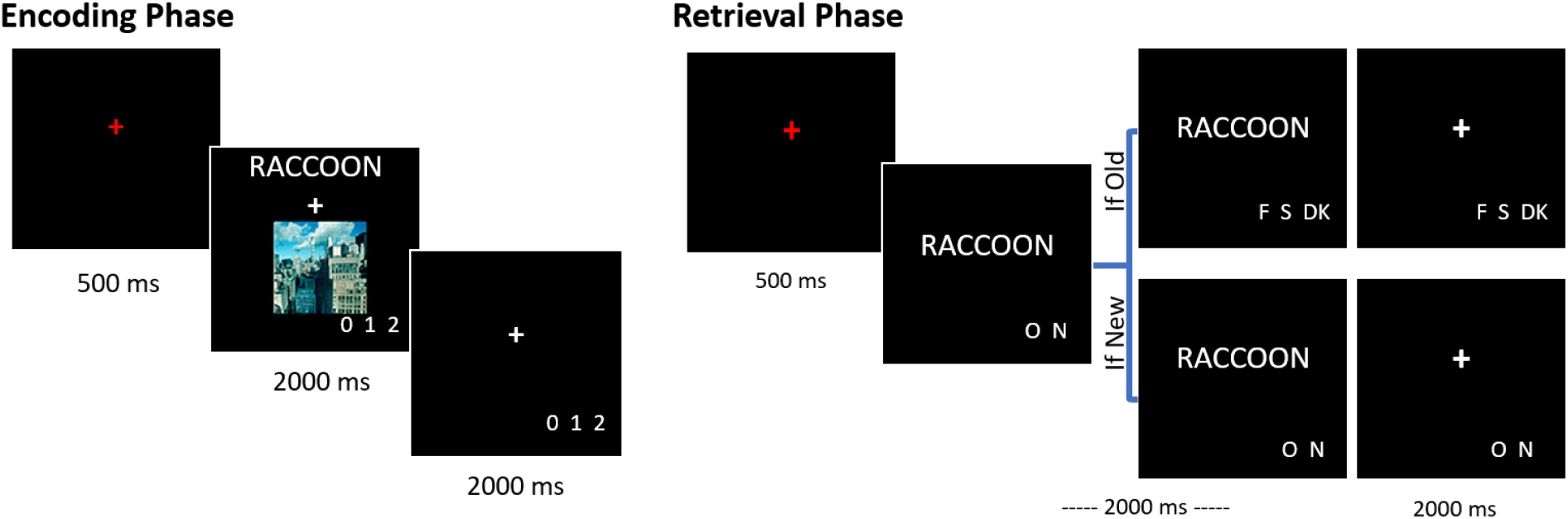
Schematic of the encoding and retrieval tasks. At encoding, participants were presented with words paired with an image of a face or a scene. At retrieval, they were presented with a test word and required to indicate whether they remembered seeing the word during the encoding phase, and if so, whether it had been paired with an image of a face or a scene.

Each test trial began with a red fixation cross for 500 ms, which was immediately replaced by the test item for a duration of 2000 ms. Response prompts appeared underneath the item at its onset. Participants were instructed first to indicate whether they remembered seeing the word at study by making an “Old” or “New” response. For each word endorsed “Old”, participants went on to make a source memory judgment, indicating whether the word had been studied with a face or a scene. A third “Don’t Know” option was included to discourage participants from guessing. As with the study phase, the test item was replaced with a white fixation cross for 2000 ms, and participants were allowed to make their memory judgments throughout the full 4000 ms response window. Responses were made using a scanner-compatible button box. Old / New responses were made with the index and middle fingers of the right hand with the ordering of the fingers counterbalanced across participants. The Face / Scene / Don’t Know responses were made with the index, middle, and ring fingers of the right hand and were also counterbalanced across participants while ensuring that the Don’t Know response was never assigned to the middle finger.

### 2.5. MRI Data Acquisition and Preprocessing

Functional and structural MRI data were acquired using a Philips Achieva 3T MRI scanner (Philips Medical System, Andover, MA, US) equipped with a 32-channel head coil. Anatomical scans were acquired with a T1-weighted 3D magnetization-prepared rapid gradient echo (MPRAGE) pulse sequence (field of view [FOV] = 256 × 256 mm, voxel size = 1 × 1 × 1 mm, 160 slices, sagittal acquisition). Functional data was obtained with a T2*-weighted echo-planar-imaging (EPI) sequence (FOV = 240 × 240 mm, TR = 2 s, TE = 30 ms, flip angle = 70°). EPI volumes consisted of 34 axial slices acquired in an ascending order parallel to the anterior-posterior commissure, with an interslice gap of 1 mm. The voxel size of the EPI volumes was 3 mm isotropic.

The MRI data were preprocessed using Statistical Parametric Mapping (SPM12, Wellcome Department of Cognitive Neurology, London, UK) and custom Matlab code (MathWorks). The functional data were realigned to the mean EPI image and slice-time corrected using *sinc* interpolation with reference to the 17^th^ slice. Following realignment, images were reoriented and normalized to a sample-specific EPI template according to previously published procedures that ensured an unbiased contribution of each age group to the template (de Chastelaine et al., 2011, 2016). Lastly, images were smoothed with an 8 mm full-width at half-maximum Gaussian kernel. The time series of the study and test runs were concatenated using the *spm_fmri_concatenate* function prior to the implementation of first-level general linear model (GLM; see below).

### 2.6. Data Analysis

#### 2.6.1. Whole-brain Univariate Analysis

The functional data were analyzed with a two-stage univariate GLM approach. At the first stage, separate GLMs were implemented for the study and test data of each participant. The study trials were binned into two events of interest (face and scene trials) and the neural activity elicited by the trials was modeled with a boxcar function extending over the 2s period during which the word-image pair remained on the screen. The boxcar regressors were convolved with two canonical hemodynamic response functions (HRFs): SPM’s canonical HRF and an orthogonalized delayed HRF. The delayed HRF was created by shifting the canonical HRF by one TR (2s) later and using the Gram-Schmidt procedure to orthogonalize it with the canonical HRF (Andrade et al., 1999). The delayed HRF did not produce any findings in addition to those described below and thus is not discussed further. In addition to the events of interest described above, the GLM for the study phase also modeled the following trials as covariates of no interest: filler trials, trials with missing or multiple responses, trials receiving a response before 500 ms or after 4500 ms following stimulus onset, and the 30-second rest period. Additional covariates of no interest comprised 6 motion regressors reflecting rigid-body translation and rotation, spike covariates regressing out volumes with transient displacement > 1 mm or > 1° in any direction, and the mean signal of each scanner run. The parameter estimates from the first level GLM were carried over to a 2 (age group: younger, older) x 2 (study trial: face, scene) mixed factorial ANOVA which was height-thresholded at p < 0.001 uncorrected, retaining only those clusters which survived FWE correction at p < 0.05.

The test phase trials were binned into five events of interest: face trials associated with a correct source memory judgement (face source correct), scene trials associated with a correct source memory judgement (scene source correct), recognized old items which received an incorrect source memory judgement or a Don’t know response (source incorrect + DK), studied items attracting an incorrect ‘new’ response (item miss), and new items attracting a correct ‘new’ response (correct rejection). Events of interest were modeled with a delta function time-locked to stimulus onset (the choice of a delta function was motivated by the presumed short-lived nature of the processing of the retrieval cue) and convolved with the canonical and orthogonalized delayed HRFs. As with the encoding data, the delayed HRF did not identify any additional clusters of interest. Covariates of no interest comprised filler trials, false alarms, trials with missing or multiple responses, trials attracting a response before 500 ms and after 4500 ms following stimulus onset, the 30 second rest periods, six motion regressors reflecting translational and rotational displacement, motion spike covariates, and the mean signal for each run. The second level GLM took the form of a 2 (age group: younger, older) x 5 (test trial: face source correct, scene source correct, source incorrect + DK, item miss, correct rejection) mixed factorial ANOVA. Analogous to the GLM of the study data, the ANOVA was height-thresholded at p < 0.001 uncorrected and clusters were retained if they exceeded the FWE corrected threshold of p < 0.05.

#### 2.6.2. Anterior shift in scene- and face-selectivity between study and test

The primary aim of the analyses described below centered on examining age differences and the functional significance of the retrieval-related anterior shift. Here, the term ‘anterior shift’ refers to a statistically significant difference in the localization of neural activity observed at encoding and retrieval, such that the retrieval of the memory of a perceptual stimulus (e.g., an image of a scene in the context of the present study) is associated with a peak response in category-selective cortex that is anterior to the peak response elicited when the image was experienced directly. The present analyses were restricted to scene- and face-selective cortical regions where significant clusters could be identified across both age groups (i.e. clusters surviving the FWE corrected threshold of p < 0.05) in both the encoding and retrieval phases (see 3.2, Whole-Brain Results). The resulting scene-selective ROIs were localized to the parahippocampal place area (PPA), medial place area (MPA; sometimes referred as retrosplenial cortex), and occipital place area (OPA). Among face-selective clusters, only the precuneus (PCU) could be identified at both encoding and retrieval. When examining the coordinates of peak scene- and face-selective responses within these regions at the individual subject level, the analyses were restricted to anatomical masks which corresponded to the cortical regions encompassing the clusters described above. Each anatomical mask was defined by reference to SPM’s Neuromorphometrics atlas with the exception of the MPA, which was not well captured by the labels provided by Neuromorphometrics and was instead defined by reference to the Atlas of Intrinsic Connectivity of Homotopic Areas (AICHA; Joliot et al., 2015). The PPA was delimited by the parahippocampal and fusiform gyrus labels. The OPA mask was created using the atlas labels for the inferior and middle occipital gyri and the PCU mask comprised the precuneus and posterior cingulate labels. The MPA was defined using the following AICHA labels: precuneus (AICHA indices: 265, 267, 269 for the left hemisphere; 266, 268, 270 for the right hemisphere), parieto-occipital (left hemisphere: 283, 285, 289, 291; right hemisphere: 284, 286, 290, 292), and posterior cingulate (left hemisphere: 253, 255; right hemisphere: 254, 256). The AICHA atlas was resampled to 3mm isotropic voxels to match the resolution of the functional data prior to ROI definition. More details about each mask are given in Table 1.

**Table 1:** Size of the anatomical regions of interest, the number of searchlights from which parameter estimates were extracted, and the mean (SD) size of the searchlights.

|  | Mask size (in voxels) | Number of searchlights | Mean (SD) searchlight size (in voxels) |
| --- | --- | --- | --- |
| **Scene ROIs** |  |  |  |
| Left PPA | 410 | 388 | 11.56 (3.14) |
| Right PPA | 404 | 381 | 11.73 (3.26) |
| Left MPA | 633 | 625 | 14.28 (3.75) |
| Right MPA | 688 | 669 | 14.51 (3.67) |
| Left OPA | 645 | 636 | 13.37 (3.55) |
| Right OPA | 561 | 550 | 13.12 (3.47) |
| **Face ROIs** |  |  |  |
| Left PCU | 913 | 902 | 13.79 (3.37) |
| Right PCU | 816 | 806 | 13.33 (3.26) |

Using the outcome of each participant’s 1^st^ level GLMs, we computed *scenes > faces* and *faces > scenes* contrasts from the encoding data, and *scene source correct > face source correct* and *face source correct > scene source correct* contrasts from the retrieval data. We performed anatomically constrained univariate searchlight analyses on spherical ROIs (i.e. searchlights) of 5 mm radius that were iteratively centered around each voxel falling within a given anatomical mask. The voxels comprising each searchlight were restricted to those that fell within the relevant mask to ensure that we did not intrude on adjacent cortical regions. This approach resulted in truncated spheres at the mask edges, and any searchlights that contained fewer than 6 voxels were eliminated from the analysis. The final numbers of searchlights included in these analyses and their sizes are given in Table 1. Note that the anatomical masks comprised voxels common to all subjects and both task phases. As a result, any task and age differences in the localization of category selectivity could not have arisen due to age or task differences in the size of the searchlights. We also conducted a secondary analysis in which the searchlights were allowed to extend over the mask edges into adjacent gray matter (as defined by SPM’s Tissue Probability Mask). This analysis yielded results that were essentially identical to those described below.

For each participant, mean across-voxel parameter estimates corresponding to the scene- and face-selective encoding and retrieval contrasts were extracted from each searchlight. We employed scene > faces (encoding) and scene source correct > face source correct (retrieval) contrasts when examining selectivity within the PPA, MPA and OPA. The face-selective face > scene and face source correct > scene source correct contrasts were used in the case of the face-selective PCU. To localize peaks manifesting maximal scene or face selectivity in each region, for each participant we ranked the searchlights in terms of their mean category-selective responses and selected the top 5%. This was done separately for the encoding and retrieval contrasts. The MNI coordinates of the centers of these spheres were then averaged across each plane to compute the coordinates of their centroid, and this defined the locus of peak selectivity. This approach resulted in two centroids for each participant and ROI, one defining the location of peak category selectivity at encoding and the other at retrieval. Encoding-retrieval shifts were defined as the distance (in mm) between the two centroids along the posterior – anterior plane (i.e. the difference between the respective Y coordinates). Thus, negative values would indicate a retrieval-related posterior shift (such that the peak category selectivity at retrieval is located posterior to the encoding peak) and positive values indicate an anterior shift (such that the retrieval peak is localized anterior to the encoding peak). A schematic describing the analysis approach is illustrated in Figure 2. To ensure the results we report below were not dependent on the choice of searchlight parameters (i.e., the searchlight radius and the number of top ranked searchlights), we conducted additional analyses employing searchlights of 3 mm, 5 mm and 8 mm radius while selecting the top 1%, 5% and 10% of searchlights to build the centroids. The effect of age group (see 3.3. *Retrieval-related Anterior Shift* below) remained stable regardless of parameter choice. A reliable relationship across participants between the size of the shift and memory performance in the PPA (see *3*.*4 Relationship with Memory Performance* below) was however evident only for the 5 mm and 8 mm searchlights. We note that since we eliminated searchlights containing fewer than 6 voxels, and a full 3 mm searchlight contained only 7 voxels, approximately 60% of the 3mm searchlights in the PPA were lost because they extended outside the boundary of the anatomical mask (by contrast, only 5% were lost in the case of the 5 mm radius searchlight). We attribute the failure to find a reliable relationship between the PPA anterior shift and memory performance when employing the 3mm searchlights to this data loss and an attendant increase in measurement noise.

**Figure 2:**
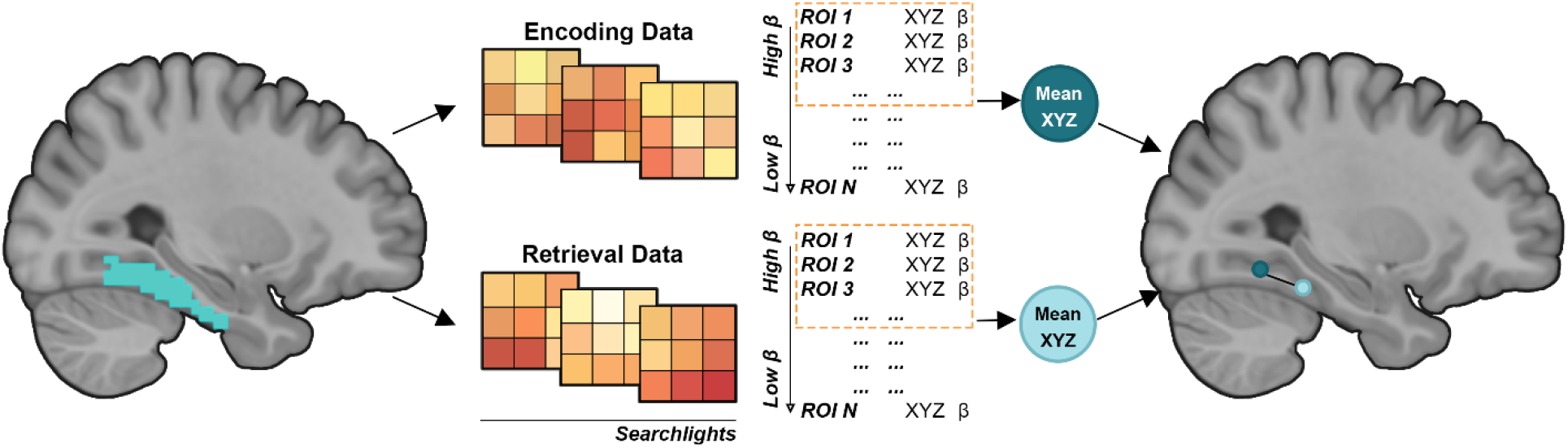
Schematic illustration of the encoding-retrieval displacement analysis pipeline. Searchlights were iteratively centered around every voxel inside a given anatomical mask. We selected the top 5% most category-selective spheres, separately for the encoding and retrieval data. The MNI coordinates of the searchlight centers were averaged to compute the center of mass (centroids) of category selectivity. The retrieval-related anterior shift was defined as the distance (in mm) between the encoding and retrieval centroids along the posterior-anterior plane. See main text for details.

#### 2.6.3. Statistical Analyses

Statistical analyses were performed using R software (R Core Team, 2020). Statistical tests were considered significant at p < 0.05 unless otherwise stated (e.g., see exploratory analyses in 3.4. *Relationship with Memory Performance*, where we correct for family-wise error). ANOVAs were performed using the afex package (Singmann et al., 2016) and degrees of freedom were corrected for nonsphericity using the Greenhouse-Geisser procedure (Greenhouse and Geisser, 1959). All t-tests were performed using the t.test function and regression analyses were performed using the lm function, both in base R. Partial correlations were conducted using pcor.test in the ppcor package (Kim, 2015).

## 3. Results

### 3.1. Behavioral Results

Behavioral performance and neuropsychological test performance have been reported previously (Srokova et al., 2020; Hill et al., 2021) and are only briefly summarized below. With regards to neuropsychological test performance, younger adults outperformed older adults on the CVLT Short Delay – Free recall, CVLT recognition – False alarms, WMS Logical Memory I and II, SDMT, Trails A and B, and Raven’s matrices. On the experimental task, item recognition (Pr) was operationalized as the difference between hit rate (the proportion of items correctly endorsed ‘old’) and the false alarm rate (the proportion of new items incorrectly endorsed ‘old’). ANOVA revealed a main effect of age (F(1,46) = 10.112, p = 0.003, partial-η^2^ = 0.180), reflective of higher Pr in younger (M [SD] = 0.68 [0.17]) relative to older adults (M [SD] = 0.54 [0.13]). The ANOVA also identified a main effect of image category (F_(1,46)_ = 5.443, p = 0.024, partial-η^2^ = 0.106), reflective of higher Pr for words studied with faces (M [SD] = 0.63 [0.16]) relative to scenes (M [SD] = 0.60 [0.15]). The interaction between age group and image category was not significant (F_(1,46)_ = 0.766, p = 0.386, partial-η^2^ = 0.016). Source memory performance (pSR) was operationalized by a single high-threshold model (Snodgrass and Corwin, 1988; see also Gottlieb et al., 2010; Mattson et al., 2014) using the formula: pSR = [pSource correct - 0.5 *(1 – pDon’t Know)] / [1 - 0.5 * (1 – pDon’t Know)], where ‘pSource Correct’ and ‘pDon’t know’ refer to the proportion of correctly recognized old trials receiving an accurate source memory judgement or a ‘Don’t Know’ response, respectively. An independent samples t-test revealed that pSR was significantly lower in older (M [SD] = 0.51 [0.16]) than in younger adults (M [SD] = 0.68 [0.18]; t_(45.51)_ = -3.440, p = 0.001).

### 3.2. Whole-Brain Results

Figure 3-A illustrates the *Scene > Face* and *Face > Scene* contrasts at encoding, and Figure 3-B depicts the *Scene source correct > Face source correct* and *Face source correct > Scene source correct* contrasts at retrieval. These results have been reported previously (Hill et al., 2021) and are re-reported here because of their relevance to the present analyses and ROI definition (we note however that Hill et al., 2021 focused on face and scene recollection contrasts [*face/scene source correct > source incorrect + DK]*, the outcomes of which are highly similar to those reported here). At encoding, scene-selective clusters were identified along the parahippocampal and fusiform gyri, extending into the retrosplenial and medial occipital cortices. Face-selective clusters were identified in the precuneus/posterior cingulate cortex, medial prefrontal cortex, and along the medial temporal lobe bilaterally extending into the amygdala and anterior hippocampus. Face-selective clusters were also evident in middle temporal gyri and the right fusiform cortex. At retrieval, scene-selective clusters were evident in bilateral parahippocampal cortex, retrosplenial cortex, and the left middle occipital cortex along with a cluster in the right orbitofrontal cortex extending into the subgenual anterior cingulate cortex. The sole face-selective cluster at retrieval was observed in the precuneus, extending into the posterior cingulate cortex.

**Figure 3:**
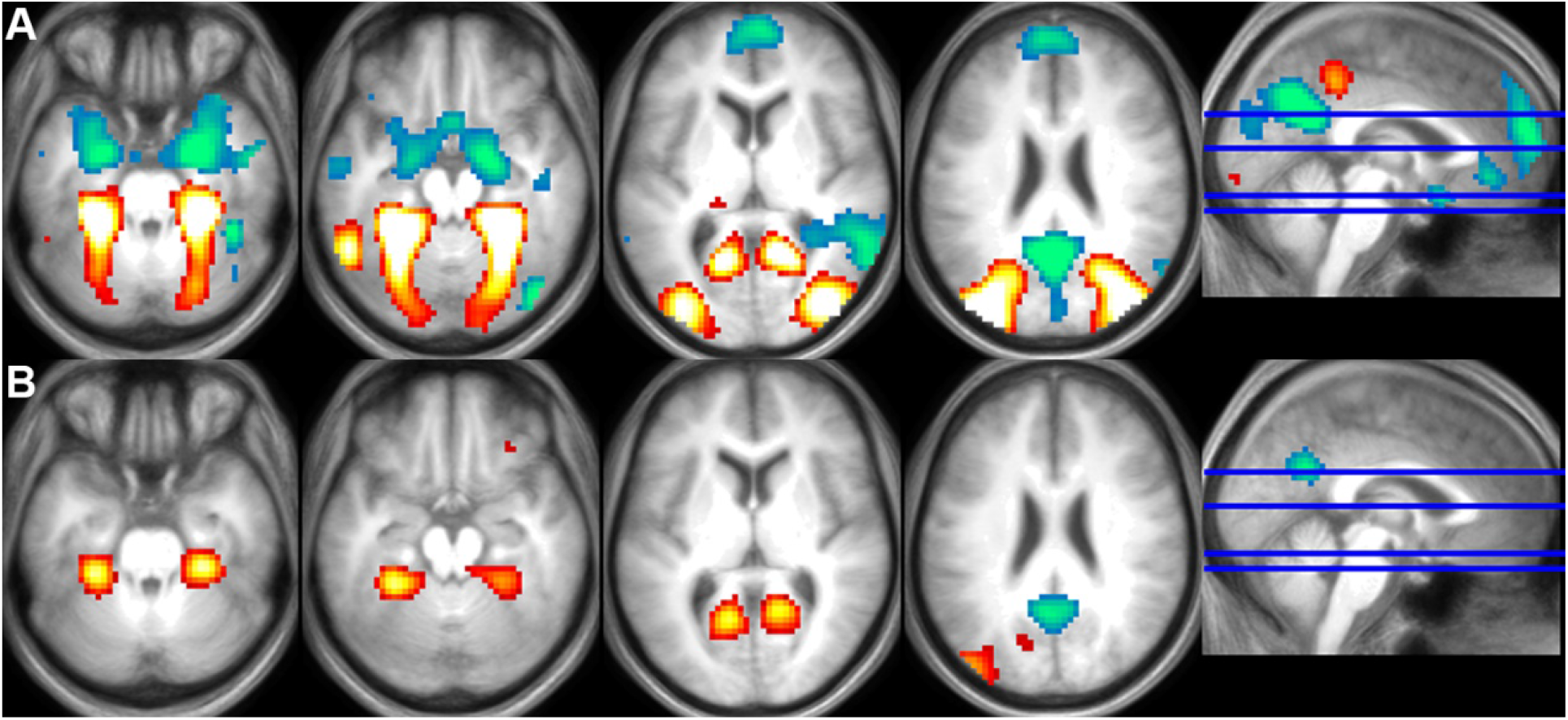
Univariate scene-selective (red) and face-selective (blue) effects at encoding **(A)** and retrieval **(B)**, collapsed across age groups. Clusters are overlaid on the across-participant mean T1 image. In both cases, clusters are displayed a p < 0.001 after FWE cluster size correction (p < 0.05).

### 3.3. Retrieval-related Anterior Shift

First, we aimed to establish which ROIs, if any, exhibited a retrieval-related anterior shift. To this end, we examined whether the coordinates of retrieval centroids were systematically displaced relative to the encoding centroids. To achieve this, for each participant we measured the distance along the Y plane in MNI space between encoding and retrieval centroids. We then tested whether these distances were significantly different from zero, using a one sample t-test. As noted previously (see 2.6.2. *Anterior shift in scene- and face-selectivity between study and test*), distance measures greater than zero indicate that the retrieval centroid is shifted anteriorly to the encoding centroid, whereas negative values indicate a posterior shift. Figure 4 depicts the encoding and retrieval centroids and their corresponding distances for each individual participant. As is evident from Table 2, in younger adults there was a reliable anterior shift in bilateral PPA and OPA, while the shift was not significantly different from zero in either the MPA (for scenes) or the PCU (for faces). By contrast, older adults exhibited a reliable anterior shift in all ROIs except for the left MPA, where it approached significance. Given the consistent trend in all ROIs towards a retrieval-related anterior shift, in the interest of clarity we refer to this simply as the ‘anterior shift’ in the analyses described below.

**Table 2:** Mean (SD) of retrieval-related anterior shift (in mm) and the outcomes of one-sample t-tests against zero.

|  | *Younger Adults* | *Older Adults* |
| --- | --- | --- |
| *Left PPA* | 7.29 (5.79)<br>t = 6.169, p < 0.001 | 11.09 (8.83)<br>t = 6.159, p < 0.001 |
| *Right PPA* | 9.03 (6.40)<br>t = 6.906, p < 0.001 | 13.82 (10.05)<br>t = 6.741, p < 0.001 |
| *Left MPA* | 1.24 (6.69)<br>t = 0.907, p = 0.374 | 3.45 (8.21)<br>t = 2.059, p = 0.051 |
| *Right MPA* | 2.25 (5.51)<br>t = 2.005, p = 0.057 | 3.70 (6.51)<br>t = 2.784, p = 0.011 |
| *Left OPA* | 3.01 (4.21)<br>t = 3.503, p = 0.002 | 4.34 (3.59)<br>t = 5.927, p < 0.001 |
| *Right OPA* | 2.83 (5.55)<br>t = 2.504, p = 0.020 | 5.65 (4.90)<br>t = 5.651, p < 0.001 |
| *Left PCU* | 0.99 (5.83)<br>t = 0.833, p = 0.413 | 5.83 (8.85)<br>t = 3.229, p = 0.004 |
| *Right PCU* | 1.71 (7.31)<br>t = 1.146, p = 0.264 | 5.33 (11.88)<br>t = 2.197, p = 0.038 |

For all t-tests: df = 23

**Figure 4.**
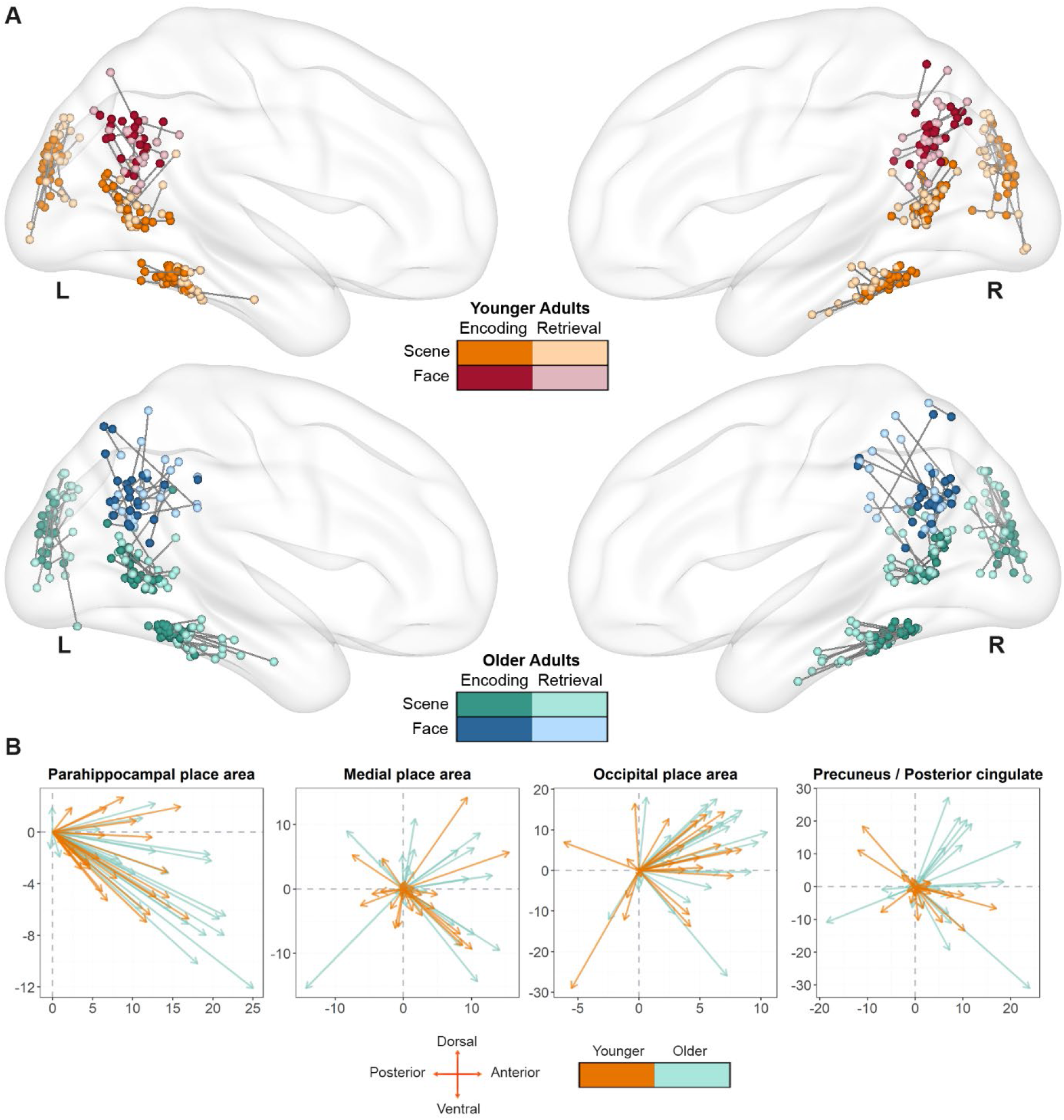
**(A)**: Encoding and Retrieval centroids for each subject plotted on a medial view of the brain surface template provided by BrainNet (Xia et al., 2013). Each subject’s centroid pair is linked with a line. **(B):** Retrieval-related shift (in mm) of the retrieval centroid (arrow) relative to the encoding centroid (origin) for each subject, collapsing across the two hemispheres.

The shift metrics were entered into a 2 (Age group) x 4 (ROI) x 2 (Hemisphere) mixed effects ANOVA. This revealed significant main effects of ROI (F_(2.25, 103.65)_ = 14.672, p < 0.001, partial-η^2^ = 0.242), and age group (F_(1,46)_ = 12.897, p = 0.001, partial-η^2^ = 0.219). The main effect of hemisphere was not significant, (F_(1,46)_ = 2.855, p = 0.098, partial-η^2^ = 0.058), and neither were any of the two- or three-way interactions (p > 0.456). The main effect of age group is indicative of a greater anterior shift in the older relative to younger adults and the absence of an ROI x age group interaction indicates that this age difference did not differ according to ROI (see Figure. 5-A). Since no hemisphere effects were identified, subsequent analyses were performed on distance measures averaged across the hemispheres.

**Figure 5.**
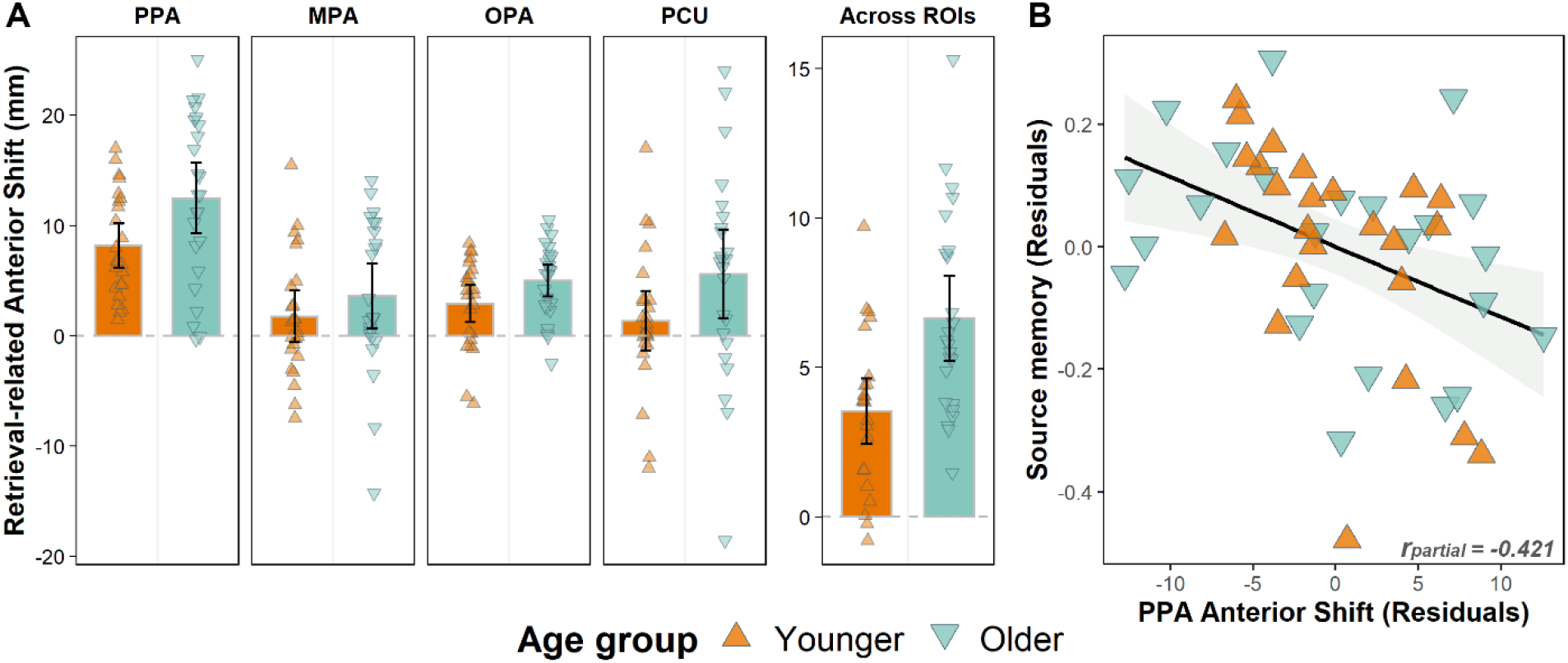
**(A):** The anterior shift plotted separately for younger and older adults in each ROI. The distance values are plotted after collapsing across hemispheres, and an additional panel is provided illustrating the distances after collapsing across all ROIs to illustrate the main effect of age. Error bars signify 95% confidence intervals. (**B):** Age-invariant relationship between retrieval-related anterior shift in the PPA and source memory performance.

### 3.4. Relationship with memory performance

We performed a series of multiple regression analyses to examine whether the retrieval-related anterior shift covaried across participants with their memory performance. Separate regression models were constructed to predict Pr (collapsed across image category) and pSR, using age group, the anterior shift in each ROI, and their interaction as predictors. The interaction term was included in the models to examine whether any relationships between the anterior shift and memory performance were moderated by age group. If the term was not significant, the regression analysis was followed-up by computing the partial correlation between the anterior shift and memory, controlling for age group. Considering that these analyses were exploratory in nature, we assessed the significance of any findings after Bonferroni correction for family-wise error (8 tests; corrected significance level: p < 0.00625). For completeness, effects that achieved significance before correction are also reported; these findings should however be interpreted with caution and are not discussed further.

The interaction term in the regression model predicting Pr from the predictors of age group and the anterior shift in the face-selective PCU was significant before (p = 0.033) but not following correction (p_(corrected)_ = 0.264). The interaction terms in the remaining regression models were not significant (p > 0.052 before correction). Thus, there was little evidence to suggest that age group moderated any potential relationships between the anterior shift and memory performance. Therefore, as noted previously, we went on to examine partial correlations between anterior shift metrics and memory performance, controlling for the influence of age group.

The only partial correlation to survive correction was that between the PPA anterior shift and pSR (r_partial_ = -0.421, p = 0.003, p_(corrected)_ = 0.024). This result (see figure 5-B) reflected the fact that, regardless of age group, greater PPA anterior shift was associated with relatively lower source memory performance. Additionally, the MPA anterior shift exhibited a sizeable negative correlation with Pr (r_partial_ = -0.377, p = 0.009), but this narrowly failed to survive correction (p_(corrected)_ = 0.072). A positive correlation between pSR and the shift in the PCU (r_partial_ = 0.316, p = 0.031) also failed to survive correction (p_(corrected)_ = 0.248). No other significant correlations were identified (p > 0.120, p_(corrected)_ = 0.960). In summary, after correcting for family-wise error, a single correlation met our criterion for statistical significance: an age-invariant correlation between the PPA anterior shift and pSR.

## 4. Discussion

In the present study, we examined age effects on the retrieval-related anterior shift and its relationship with memory performance. In both young and older adults, we identified robust evidence for an anterior shift in two scene-selective cortical regions (PPA & OPA). In addition, in older adults only, the shift was reliable for scenes in the MPA and for faces in the PCU. Of importance, independently of ROI, the anterior shift was robustly larger in the older group. Moreover, the magnitude of the shift in the scene-selective PPA demonstrated an age-invariant, negative correlation with source memory performance. In sum, consistent with the predictions outlined in the Introduction, the retrieval-related anterior shift covaries positively with increasing age and negatively with memory performance.

The earliest findings suggestive of systematic differences in the loci of cortical activity associated with perception versus memory were reported in studies contrasting color perception and color imagery (e.g., Chao & Martin, 1999; Simmons et al., 2007). Extending these findings, more recent research has reported that regions recruited during scene retrieval and scene imagery are localized anteriorly to the regions recruited during scene perception (e.g., Chrastil 2018; Silson et al., 2019; Bainbridge et al., 2021; Steel et al., 2021). Such findings motivated the proposal that scene-selective cortical regions can be sub-divided into two networks (Baldassano et al., 2016). The ‘posterior scene network’ is held to include retinotopically organized regions responsible for processing visual input, while the ‘anterior network’, which includes the hippocampus as one of its constituents, supports scene representations retrieved from memory as well as spatial navigation and other memory-guided behaviors (e.g., visual exploration). This subdivision is held to be honored within the PPA, which has been partitioned into posterior (perceptual) and anterior (mnemonic) sub-divisions (Baldassano et al., 2016). However, our findings, which demonstrate that the size of the anterior shift is sensitive to age and memory performance, challenge the view that regions such as the PPA can be dichotomized into functionally distinct posterior and anterior sub-regions. Furthermore, findings of analogous shifts in other scene-selective cortical regions such as the OPA (see Figure 5-A), and for other perceptual categories (Lee et al. 2012), add further weight to the proposal that the anterior shift reflects more than a segregation between two functional networks (see also the discussion of the present findings for faces below).

In a recent review, Favila and colleagues (2020) proposed that mnemonic representations undergo a ‘transformation’, with perceived and retrieved representations differing in quality, content and amount of information. This transformation reflects a differential weighting of episodic attributes, such that a retrieved representation is biased towards the representation of ‘high-level’ conceptual information at the expense of lower-level perceptual detail. The differential emphasis on higher-versus lower-level features is reflected in the localization of concurrent cortical activity. It has been conjectured that cortical regions extending along the posterior-anterior axis are hierarchically organized, such that more posterior regions support the processing of relatively low level stimulus properties, while more anterior regions support higher-level semantic or conceptual processing (e.g. Simmons et al., 2007, Peelen & Caramazza, 2012). This proposal implies the existence of a processing gradient along which modality-specific perceptual properties are increasingly ‘abstracted away’ at the expense of higher-level conceptual features (see Introduction). This ‘abstraction account’ leads to the prediction that the magnitude of the anterior shift will vary depending on whether a retrieval test requires retrieval primarily of conceptual information as opposed to high-fidelity, modality-specific detail (Simmons et al., 2007).

To date, reports of the anterior shift have been confined to young adults. The data from the present study extend these findings by demonstrating that the shift is exaggerated in older adults. As noted above, the abstraction account of the shift implies that it reflects a representational ‘transformation’ that de-emphasizes perceptual detail. This account allows for a simple explanation of the present effects of age on the anterior shift, given the extensive evidence that retrieved episodic information contains less detail, and is more ‘gist-based’ in older than younger adults (Koustaal & Schacter, 1997; Dennis et al., 2007; 2008, Gallo et al., 2019). That is, whereas memory for the gist of an event is relatively spared in older adults, the retention of more fine-grained, individuating features of an episode appears to be especially susceptible to increasing age (Nilakantan et al., 2018; Korkki et al., 2020). We propose that the neural expression of this age difference in retrieved episodic content accounts for the exaggerated anterior shifts evident in our older sample in the present study.

Of importance, we identified a reliable anterior shift not only in scene selective cortical regions, but in the face selective PCU also, albeit in older adults only. Our failure to identify an anterior shift for faces in the PCU in younger adults is consistent with prior findings that face stimuli do not elicit a retrieval-related anterior shift in this population (Steel et al., 2021). There are two possible explanations for why we find an anterior shift in the PCU for faces in older but not younger adults. First, it could be that this effect is specific to older adults; that is, for unknown reasons, younger adults did not retrieve face representations that were abstracted away from the original stimulus event. Alternatively, the seeming absence of a PCU effect in the younger adults might merely be a consequence of the fact that the shift in this region is smaller than that in other regions (compare, for example, the magnitude of the shift in the PPA vs. the PCU in the older participants illustrated in figure 5-A). By this argument, a shift might be detectable in the PCU of young adults given sufficient spatial resolution and statistical power.

Whereas the notion of a cortical posterior-anterior gradient from perception to memory is well supported, the question whether the magnitude of the retrieval-related anterior shift impacts memory performance has been largely unexplored (but see Davis et al., 2021). Here, we sought for relationships between the anterior shift and memory performance on the assumption that memory for the details of an event is more likely to be accurate when there is strong overlap (indexed by a relatively small anterior shift) between experienced and retrieved event representations (see Introduction). Consistent with this prediction, we identified a negative, age-invariant correlation between the PPA anterior shift and source memory performance. That is, regardless of age group, a greater anterior shift was associated with poorer memory for the study pairs. This finding supports the proposal that the localization of retrieval-related neural activity has implications for the content of retrieval, and it is also consistent with the notion that more anterior regions of the PPA support mnemonic representations containing relatively sparse perceptual detail (Bainbridge et al., 2021; Steel et al., 2021). As alluded to earlier, the finding that the shift (at least, in the PPA) both correlates with memory performance and is enhanced in older adults suggests an intriguing mechanism that might partially account for age-related memory decline.

In conclusion, the present study revealed robust age differences in the retrieval-related anterior shift in both scene-and face-selective cortical regions. We also demonstrate that the shift is (negatively) associated with source memory performance, supporting the notion that low- and high-level stimulus information is represented in different cortical regions at multiple levels of abstraction along the posterior-anterior axis. Future research should examine whether the age effects observed here extend to other stimulus categories (such as objects) or other modalities (e.g. auditory stimuli). In sum, the findings reported here shed light on the functional significance of the anterior shift in relation to memory accuracy and potentially provide an increased understanding of the factors contributing to age-related memory decline.

## Acknowledgements

This work was supported by the National Science Foundation Grant 1633873 and the National Institute of Aging Grant RF1AG039103.

## Notes

**Conflicts of Interest:** None

### Competing Interest Statement

The authors have declared no competing interest.

## References

Addis, D. R., Wong, A. T., & Schacter, D. L. (2008). Age-related changes in the episodic simulation of future events. Psychological Science, 19(1), 33–41. https://doi.org/10.1111/j.1467-9280.2008.02043.x

Andrade, A., Paradis, A.-L., Rouquette, S., & Poline, J.-B. (1999). Ambiguous Results in Functional Neuroimaging Data Analysis Due to Covariate Correlation. NeuroImage, 10(4), 483–486. https://doi.org/10.1006/nimg.1999.0479

Baldassano, C., Esteva, A., Fei-Fei, L., & Beck, D. M. (2016). Two Distinct Scene-Processing Networks Connecting Vision and Memory. ENeuro, 3(5). https://doi.org/10.1523/ENEURO.0178-16.2016

Bowman, C. R., Chamberlain, J. D., & Dennis, N. A. (2019a). Sensory Representations Supporting Memory Specificity: Age Effects on Behavioral and Neural Discriminability. The Journal of Neuroscience, 39(12), 2265–2275. https://doi.org/10.1523/JNEUROSCI.2022-18.2019

Bowman, C. R., Chamberlain, J. D., & Dennis, N. A. (2019b). Sensory Representations Supporting Memory Specificity: Age Effects on Behavioral and Neural Discriminability. Journal of Neuroscience, 39(12), 2265–2275. https://doi.org/10.1523/JNEUROSCI.2022-18.2019

Chao, L. L., & Martin, A. (1999). Cortical Regions Associated with Perceiving, Naming, and Knowing about Colors. Journal of Cognitive Neuroscience, 11(1), 25–35. https://doi.org/10.1162/089892999563229

Chrastil, E. R. (2018). Heterogeneity in human retrosplenial cortex: A review of function and connectivity. Behavioral Neuroscience, 132(5), 317–338. https://doi.org/10.1037/bne0000261

Danker, J. F., & Anderson, J. R. (2010). The ghosts of brain states past: Remembering reactivates the brain regions engaged during encoding. Psychological Bulletin, 136(1), 87–102. https://doi.org/10.1037/a0017937

Davis, S. W., Geib, B. R., Wing, E. A., Wang, W.-C., Hovhannisyan, M., Monge, Z. A., & Cabeza, R. (2021). Visual and Semantic Representations Predict Subsequent Memory in Perceptual and Conceptual Memory Tests. Cerebral Cortex, 31(2), 974–992. https://doi.org/10.1093/cercor/bhaa269

de Chastelaine, M., Mattson, J. T., Wang, T. H., Donley, B. E., & Rugg, M. D. (2016). The relationships between age, associative memory performance, and the neural correlates of successful associative memory encoding. Neurobiology of Aging, 42, 163–176. https://doi.org/10.1016/j.neurobiolaging.2016.03.015

de Chastelaine, M., Wang, T. H., Minton, B., Muftuler, L. T., & Rugg, M. D. (2011). The Effects of Age, Memory Performance, and Callosal Integrity on the Neural Correlates of Successful Associative Encoding. Cerebral Cortex, 21(9), 2166–2176. https://doi.org/10.1093/cercor/bhq294

Delis DC, Kramer JH, Kaplan E, Ober BA (2000) California verbal learning test, Ed 2. San Antonio: The Psychological Corporation.

Dennis, N. A., Kim, H., & Cabeza, R. (2007). Effects of aging on true and false memory formation: An fMRI study. Neuropsychologia, 45(14), 3157–3166. https://doi.org/10.1016/j.neuropsychologia.2007.07.003

Dennis, N. A., Kim, H., & Cabeza, R. (2008). Age-related differences in brain activity during true and false memory retrieval. Journal of Cognitive Neuroscience, 20(8), 1390–1402. https://doi.org/10.1162/jocn.2008.20096

Favila, S. E., Lee, H., & Kuhl, B. A. (2020). Transforming the Concept of Memory Reactivation. Trends in Neurosciences, 43(12), 939–950. https://doi.org/10.1016/j.tins.2020.09.006

Folville, A., Bahri, M. A., Delhaye, E., Salmon, E., D’Argembeau, A., & Bastin, C. (2020). Age-related differences in the neural correlates of vivid remembering. NeuroImage, 206, 116336. https://doi.org/10.1016/j.neuroimage.2019.116336

Gallo, H. B., Hargis, M. B., & Castel, A. D. (2019). Memory for Weather Information in Younger and Older Adults: Tests of Verbatim and Gist Memory. Experimental Aging Research, 45(3), 252–265. https://doi.org/10.1080/0361073X.2019.1609163

Greenhouse, S. W., & Geisser, S. (1959). On methods in the analysis of profile data. Psychometrika, 24(2), 95–112. https://doi.org/10.1007/BF02289823

Hill, P. F., King, D. R., & Rugg, M. D. (2021). Age Differences In Retrieval-Related Reinstatement Reflect Age-Related Dedifferentiation At Encoding. Cerebral Cortex, 31(1), 106–122. https://doi.org/10.1093/cercor/bhaa210

Johnson, J. D., McDuff, S. G. R., Rugg, M. D., & Norman, K. A. (2009). Recollection, Familiarity, and Cortical Reinstatement: A Multivoxel Pattern Analysis. Neuron, 63(5), 697–708. https://doi.org/10.1016/j.neuron.2009.08.011

Joliot, M., Jobard, G., Naveau, M., Delcroix, N., Petit, L., Zago, L., Crivello, F., Mellet, E., Mazoyer, B., & Tzourio-Mazoyer, N. (2015). AICHA: An atlas of intrinsic connectivity of homotopic areas. Journal of Neuroscience Methods, 254, 46–59. https://doi.org/10.1016/j.jneumeth.2015.07.013

Kim S. (2015) ppcor: an R package for a fast calculation to semi-partial correlation coefficients. Commun Stat Appl Methods 22:665–674.

Korkki, S. M., Richter, F. R., Jeyarathnarajah, P., & Simons, J. S. (2020). Healthy ageing reduces the precision of episodic memory retrieval. Psychology and Aging, 35(1), 124–142. https://doi.org/10.1037/pag0000432

Koutstaal, W., & Schacter, D. L. (1997). Gist-based false recognition of pictures in older and younger adults. Journal of Memory and Language, 37(4), 555–583. https://doi.org/10.1006/jmla.1997.2529

Lee, S.-H., Kravitz, D. J., & Baker, C. I. (2012). Disentangling visual imagery and perception of real-world objects. Neuroimage, 59(4), 4064–4073. https://doi.org/10.1016/j.neuroimage.2011.10.055

Levine, B., Svoboda, E., Hay, J. F., Winocur, G., & Moscovitch, M. (2002). Aging and autobiographical memory: Dissociating episodic from semantic retrieval. Psychology and Aging, 17(4), 677–689.

Martin, C. B., Douglas, D., Newsome, R. N., Man, L. L., & Barense, M. D. (2018). Integrative and distinctive coding of visual and conceptual object features in the ventral visual stream. ELife, 7, e31873. https://doi.org/10.7554/eLife.31873

Nilakantan, A. S., Bridge, D. J., VanHaerents, S., & Voss, J. L. (2018). Distinguishing the precision of spatial recollection from its success: Evidence from healthy aging and unilateral mesial temporal lobe resection. Neuropsychologia, 119, 101–106. https://doi.org/10.1016/j.neuropsychologia.2018.07.035

Nilsson, L.-G. (2003). Memory function in normal aging. Acta Neurologica Scandinavica, 107(s179), 7– 13. https://doi.org/10.1034/j.1600-0404.107.s179.5.x

Nyberg, L., Lövdén, M., Riklund, K., Lindenberger, U., & Bäckman, L. (2012). Memory aging and brain maintenance. Trends in Cognitive Sciences, 16(5), 292–305. https://doi.org/10.1016/j.tics.2012.04.005

Peelen, M. V., & Caramazza, A. (2012). Conceptual Object Representations in Human Anterior Temporal Cortex. Journal of Neuroscience, 32(45), 15728–15736. https://doi.org/10.1523/JNEUROSCI.1953-12.2012

Rissman, J., & Wagner, A. D. (2012). Distributed representations in memory: Insights from functional brain imaging. Annual Review of Psychology, 63, 101–128. https://doi.org/10.1146/annurev-psych-120710-100344

R Core Team (2017) R: a language and environment for statistical computing. Vienna: R Foundation

Raven J., Raven J.C., Courth J.H. (2000) The advanced progressive matrices. In: Manual for Raven’s progressive matrices and vocabulary scales, Section 4. San Antonio: Harcourt Assessment

Reitan R.M., Wolfson D (1985) The Halstead-Reitan neuropsychological test battery: therapy and clinical interpretation. Tucson: Neuropsychological.

Rugg, M. D., Johnson, J. D., & Uncapher, M. R. (2015). Encoding and retrieval in episodic memory: Insights from fMRI. In The Wiley handbook on the cognitive neuroscience of memory (pp. 84–107). Wiley Blackwell. https://doi.org/10.1002/9781118332634.ch5

Rugg, M. D., & Thompson-Schill, S. L. (2013). Moving Forward With fMRI Data. Perspectives on Psychological Science, 8(1), 84–87. https://doi.org/10.1177/1745691612469030

Silson, E. H., Steel, A., Kidder, A., Gilmore, A. W., & Baker, C. I. (2019). Distinct subdivisions of human medial parietal cortex support recollection of people and places. ELife, 8, e47391. https://doi.org/10.7554/eLife.47391

Simmons, W. K., Ramjee, V., Beauchamp, M. S., McRae, K., Martin, A., & Barsalou, L. W. (2007). A common neural substrate for perceiving and knowing about color. Neuropsychologia, 45(12), 2802– 2810. https://doi.org/10.1016/j.neuropsychologia.2007.05.002

Singmann H., Bolker B., Westfall J., Aust F. (2016) afex: analysis of factorial experiments. Vienna: R Foundation.

Smith A. (1982) Symbol digit modalities test (SDMT) manual. Los Angeles: Western Psychological Services.

Spreen O., Benton A.L. (1977) Neurosensory center comprehensive examination for aphasia. Victoria: Neuropsychology Laboratory

Srokova, S., Hill, P. F., Koen, J. D., King, D. R., & Rugg, M. D. (2020). Neural Differentiation is Moderated by Age in Scene-Selective, But Not Face-Selective, Cortical Regions. ENeuro, 7(3). https://doi.org/10.1523/ENEURO.0142-20.2020

Steel, A., Billings, M. M., Silson, E. H., & Robertson, C. E. (2021). A network linking scene perception and spatial memory systems in posterior cerebral cortex. Nature Communications, 12(1), 2632. https://doi.org/10.1038/s41467-021-22848-z

St-Laurent, M., & Buchsbaum, B. R. (2019). How Multiple Retrievals Affect Neural Reactivation in Young and Older Adults. The Journals of Gerontology: Series B, 74(7), 1086–1100. https://doi.org/10.1093/geronb/gbz075

Trelle, A. N., Carr, V. A., Guerin, S. A., Thieu, M. K., Jayakumar, M., Guo, W., Nadiadwala, A., Corso, N. K., Hunt, M. P., Litovsky, C. P., Tanner, N. J., Deutsch, G. K., Bernstein, J. D., Harrison, M. B., Khazenzon, A. M., Jiang, J., Sha, S. J., Fredericks, C. A., Rutt, B. K., … Wagner, A. D. (2020). Hippocampal and cortical mechanisms at retrieval explain variability in episodic remembering in older adults. ELife, 9, e55335. https://doi.org/10.7554/eLife.55335

Wang, T. H., Johnson, J. D., de Chastelaine, M., Donley, B. E., & Rugg, M. D. (2016). The Effects of Age on the Neural Correlates of Recollection Success, Recollection-Related Cortical Reinstatement, and Post-Retrieval Monitoring. Cerebral Cortex, 26(4), 1698–1714. https://doi.org/10.1093/cercor/bhu333

Wechsler D. (1981) WAIS-R: Wechsler adult intelligence scale-revised. New York: The Psychological Corporation.

Wechsler D. (2001) Wechsler test of adult reading. San Antonio: The Psychological Corporation.

Wechsler D. (2009) Wechsler memory scale, Ed 4. San Antonio: The Psychological Corporation.

Xia, M., Wang, J., & He, Y. (2013). BrainNet Viewer: A Network Visualization Tool for Human Brain Connectomics. PLOS ONE, 8(7), e68910. https://doi.org/10.1371/journal.pone.0068910

Xue, G. (2018). The Neural Representations Underlying Human Episodic Memory. Trends in Cognitive Sciences, 22(6), 544–561. https://doi.org/10.1016/j.tics.2018.03.004

